# RNA-Seq in 296 phased trios provides a high resolution map of genomic imprinting

**DOI:** 10.1101/269449

**Authors:** Bharati Jadhav, Ramin Monajemi, Kristina K. Gagalova, Daniel Ho, Harmen H.M. Draisma, Mark A. van de Wiel, Lude Franke, Bastiaan T. Heijmans, Joyce van Meurs, Rick Jansen, GoNL Consortium, BIOS Consortium, Peter A.C. ‘t Hoen, Andrew J. Sharp, Szymon M. Kiełbasa

## Abstract

Combining allelic analysis of RNA-Seq data with phased genotypes in family trios provides a powerful method to detect parent-of-origin biases in gene expression. We report findings in 296 family trios from two large studies: 165 lymphoblastoid cell lines from the 1000 Genomes Project, and 131 blood samples from the Genome of the Netherlands participants (GoNL). Based on parental haplotypes we identified >2.8 million transcribed heterozygous SNVs phased for parental origin, and developed a robust statistical framework for measuring allelic expression. We identified a total of 45 imprinted genes and one imprinted unannotated transcript, 17 of which have not previously been reported as showing parental expression bias. Multiple novel imprinted transcripts showing incomplete parental expression bias were located adjacent to known strongly imprinted genes. For example, *PXDC1*, a gene which lies adjacent to the paternally-expressed gene *FAM50B*, shows a 2:1 paternal expression bias. Other novel imprinted genes had promoter regions that coincide with sites of parentally-biased DNA methylation identified in blood from uniparental disomy (UPD) samples, thus providing independent validation of our results. Using the stranded nature of the RNA-Seq data in LCLs we identified multiple loci with overlapping sense/antisense transcripts, of which one is expressed paternally and the other maternally. Using a sliding window approach, we searched for imprinted expression across the entire genome, identifying a novel imprinted putative lncRNA in 13q21.2. Our methods and data provide a robust and high resolution map of imprinted gene expression in the human genome.

## Introduction

Genomic imprinting is a special case of mono-allelic expression where genes are expressed in a parent-of-origin (PofO) specific manner. This type of mono-allelic expression can be observed in mammals at different developmental stages and is dependent on stage, cell and tissue type. Genomic imprinting plays a vital role in normal development, and errors of imprinting can underlie developmental disorders and contribute to certain cancers (Moore and Oakey 2011). Imprinting significantly influences the development of cell lineages, prenatal growth, normal brain function and metabolism (Bartolomei and Ferguson-Smith 2011). Any disruption to imprinted genes can lead to disturbed gene function and can have a deleterious effect on health. If such disruption happens at imprinted loci, it can result in imprinting disorder such as Beckwith-Wiedemann, Silver-Russell (Azzi et al. 2009), Prader-Willi, Angelman syndromes (Nicholls, Saitoh, and Horsthemke 1998), neonatal diabetes (Mackay et al. 2008) and cancer. Wilm’s tumor, colorectal cancer, and hepatoblastoma are few examples of cancer caused due to aberrant imprinting in *IGF2* gene (Steenman et al. 1994; Kaneda and Feinberg 2005).

There are many screening methods developed and applied to discover imprinted genes such as DNA methylation, histone modification, and gene expression assays. RNA sequencing (RNA-Seq) is the most direct and comprehensive way to identify imprinted genes as it allows for quantifying relative expression of the maternal and paternal alleles (allele specific expression or ASE) at all heterozygous sites with sufficient coverage. However, the technology is subject to several technical biases resulting in potential false positives (Piskol, Ramaswami, and Li 2013). The reference bias, caused by additional penalties in the alignment for non-reference alleles is the most prominent of these biases (Castel et al. 2015). Moreover, the availability of additional DNA genotype information is essential because the heterozygous sites may appear as homozygous in the RNA because of mono-allelic expression of the imprinted genes. Typically, such studies are performed without allelic inheritance information and make use of the bimodal distribution of the expression at heterozygous sites. This type of analyses lacks the ability to identify directionality of parental bias (*i.e.* assessing maternal versus paternal imprinting). Adding PofO information allows determination of maternal vs. paternal allele-specific expression with more power, in particular in the case of incomplete imprinting (slight bias towards the paternal or maternal allele), where bimodality in the distribution is difficult to assess. The use of PofO information is straightforward in mouse studies where reciprocal cross design is often used to identify maternal/paternal gene expression and imprinted genes (Gregg et al. 2010; Wang et al. 2008). So far, there are few studies performed in humans where PofO information is available. However, those studies are usually limited to either small number of trios or analysis at specific loci (Baran et al. 2015; Morcos et al. 2011)(Metsalu et al. 2014; Apostolidou et al. 2007).

To circumvent these limitations we present a robust genome-wide approach to find PofO specific gene expression and identify the signature of imprinted genes at heterozygous sites using phased DNA genotypes from parent-offspring trios and RNA-Seq data aggregated at gene level. Our method is applied to two large scale studies with a total of 296 trios: 165 trios from the HapMap / 1000 genomes projects with RNA-Seq data from lymphoblastoid cell lines (LCLs), and 131 trios from the Genome-of-the-Netherlands (Genome of the Netherlands Consortium 2014). We focus on the identification of genes and transcripts that are consistently imprinted in the population, detecting both complete imprinting (exclusive expression of the paternal or maternal allele) or incomplete imprinting (bias in expression towards the maternal or paternal allele).

## Results

We tested for imprinted gene expression using allele-specific RNA-Seq analysis of 296 parent-offspring trios derived from two independent cohorts: (i) 165 LCLs collected as part of the HapMap project, and (ii) 131 whole blood (WB) samples studied by the GoNL Consortium. In each cohort, we used phased genotypes to compute the relative expression from the maternal and paternal alleles in RNA-Seq reads at expressed heterozygous single nucleotide variants (SNVs). We analyzed 23,003 Gencode genes which had at least one heterozygous SNV with ≥1 overlapping RNA-Seq reads in >10% of the samples, and summed the paternal and maternal counts for all heterozygous SNVs contained in a gene, irrespective of their exonic or intronic nature. The inclusion of intronic SNVs increased the power of our test considerably despite their low individual coverage, as there were generally many more informative intronic than exonic SNVs. We applied two statistical tests to check for consistent parental expression bias of autosomal genes within the populations. The rationale for using two statistical tests, Wilcoxon Signed Rank (WSR) test and ShrinkBayes (SB), is their differences in power and false positive rate in case of low numbers of informative individuals and low expression. More details are given in the Supplementary Note.

Quantile-Quantile plots showed a clear excess of genes with highly significant observed p-values above the null expectation with both statistical tests and cohorts, indicating strong evidence for imprinting. Furthermore, there was no evidence of genomic inflation in our study, with all values of λ between 0.9999 and 1.02 (Figure 1). To increase resolution and avoid confounding in cases where multiple different transcripts overlapped each other, we used Unique Gene Fragments (UGF’s, see methods) annotation as the basic genomic units. Overall a total of 78 UGFs across the two populations showed significant evidence of imprinting (Supplementary Tables 1 and 2): 66 in LCLs and 43 in WB. However, the presence of overlapping transcripts, some of which were split into multiple separate annotations by our use of UGFs, created redundancy in this list. After removal of these redundancies, we further manually curated signals to (i) assign signals of imprinted expression to the gene annotation which showed best consistency with the strand and location of data, (ii) remove transcripts where biased expression was driven by outlier samples with extreme read depth, and (iii) at loci containing multiple overlapping gene annotations, to avoid inflating the number of reported genes, we removed anonymous transcripts which appeared to represent partial gene fragments (see comments in Supplementary Table 1). This identified a total of 45 imprinted genes across the two cohorts: 38 in LCLs and 31 in WB, with 23 identified in both populations (Figure 1). The paternal ratios for each of these genes in each individual are plotted in Figure 2. Among the list of 45 genes, 29 genes have been previously reported as imprinted, while 16 are putative novel imprinted genes (Tables 1 and 2).

**Figure 1.**
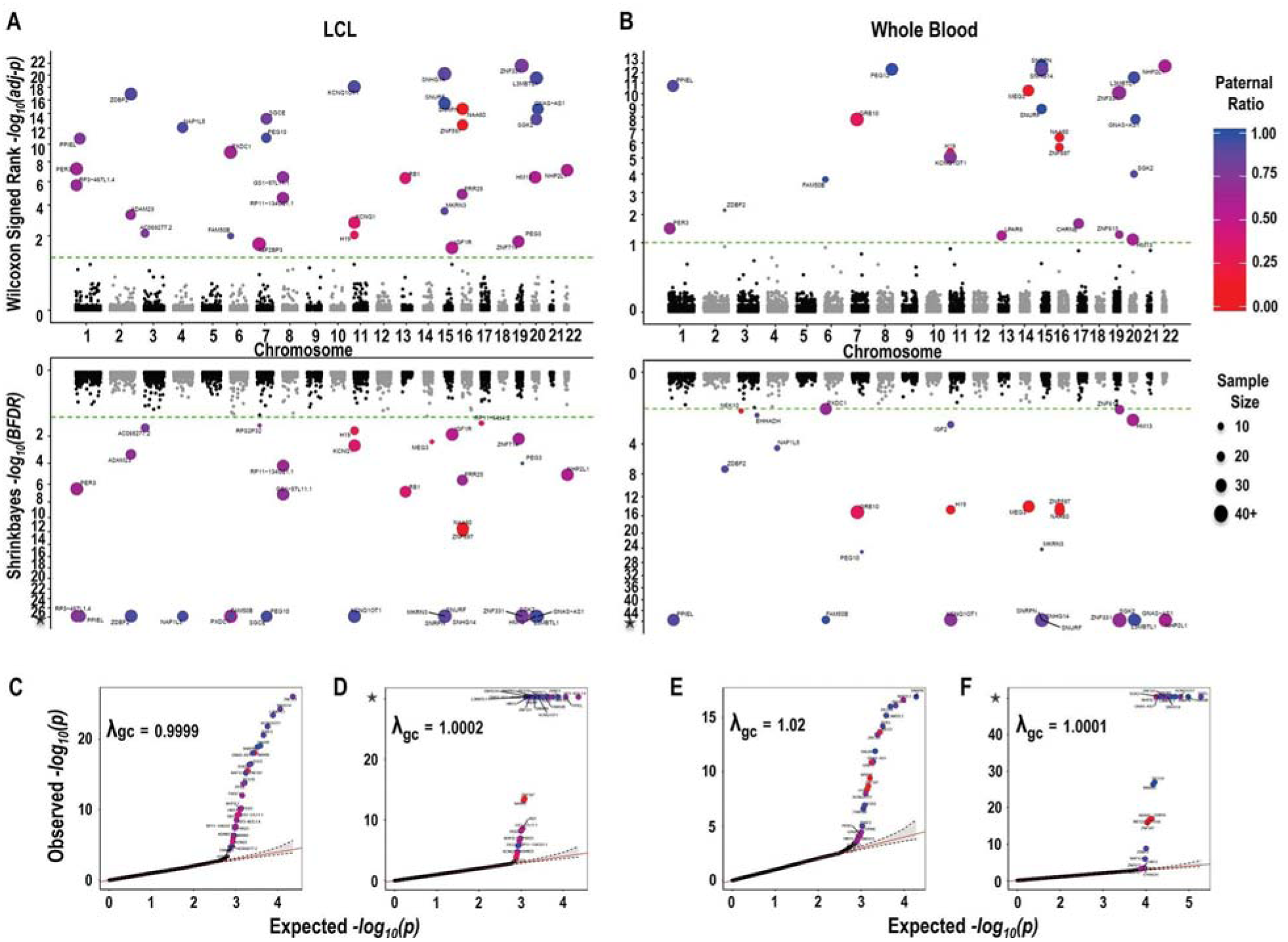
Miami and Quantile-Quantile plots of genome-wide results for parentally biased gene expression in 165 lymphoblastoid cell lines (LCL) and 131 whole blood (WB) samples. All data shown are based on bidirectional RNA-Seq data. In both **(A)** LCLs and **(B)** whole blood two statistical tests for parental bias were used: the upper panel in each cohort shows results from the paired Wilcoxon Signed Rank test, and the lower panel shows results from the *Shrinkbayes* test. −log_10_ transformed adjusted p-values are shown on the y-axis, and chromosome and position on the x-axis: the dotted green lines indicate a statistical threshold of 10% FDR, with all genes exceeding this highlighted and labeled according to their paternal expression ratio, and number of informative samples (see legend). These plots show the results of analysis based on known transcript annotations, and thus do not include the novel unannotated transcript at 13q21.2 identified by sliding window analysis. **(C and E)** QQ plots for the paired Wilcoxon Signed Rank test in LCLs and whole blood. **(D and F)** QQ plots for *Shrinkbayes* in LCLs and whole blood. Note for *Shrinkbayes*, some of the observed –log_10_ p-values are infinite, indicated by an asterisk on the y-axis. In each plot, the top 30 genes are highlighted and colored according to their paternal ratio. For both cell cohorts and statistical tests the genomic inflation factor is approximately equal to 1. For genes with multiple UGFs (Supplementary Table 2), we only plot data for the UGF with the most significant p-value.

**Figure 2.**
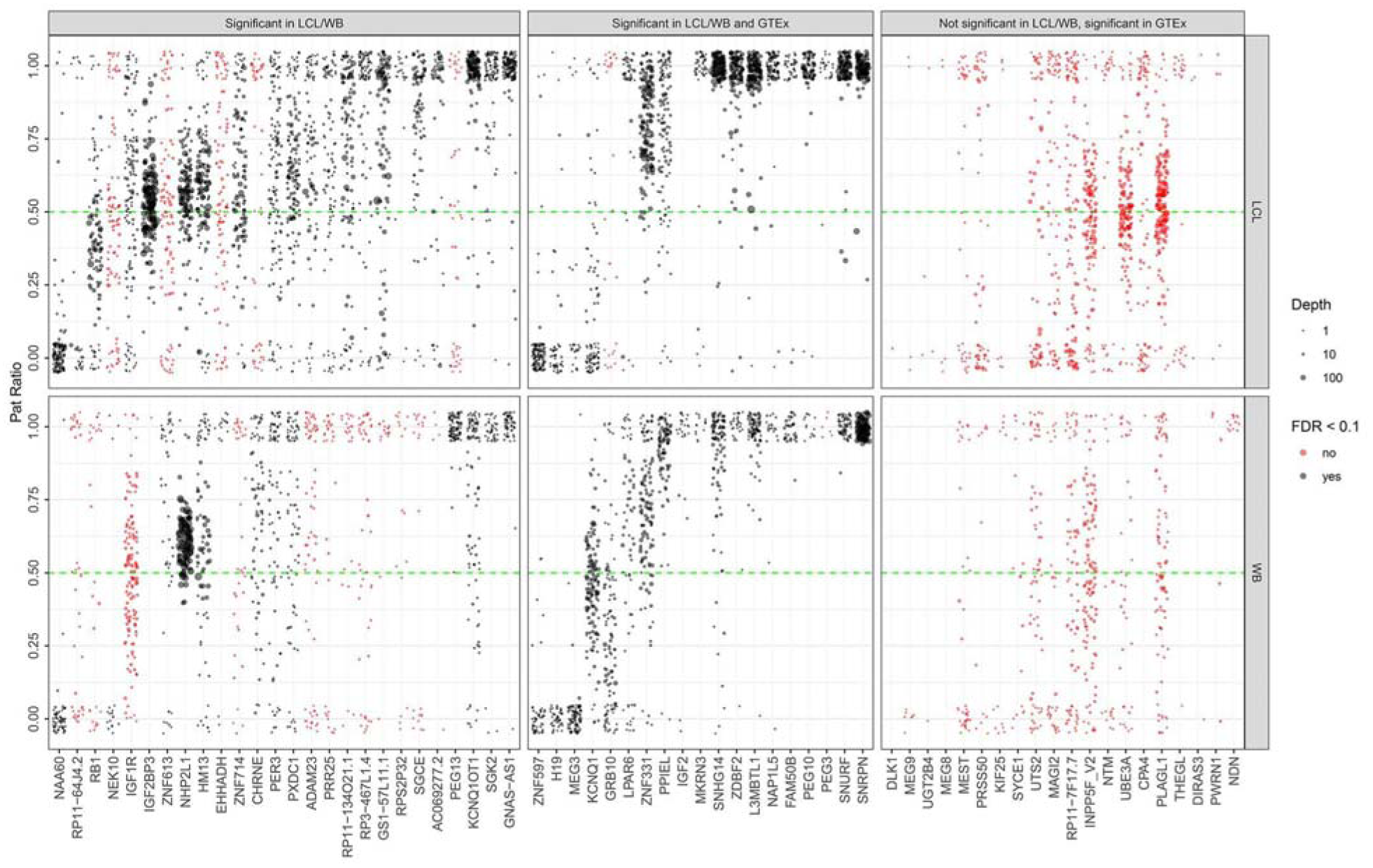
Varying degrees of parental bias among imprinted genes detected in LCLs, WB and GTEx. Each point represents the PatRatio (the mean fraction of reads transcribed from the paternal allele) in each informative individual per gene, with the point size indicating total read depth over all heterozygous transcribed SNVs in that sample. Genes are ordered left to right by increasing mean PatRatio. The upper panel shows stranded data from LCLs, while the lower panel shows unstranded data from WB samples. Note that due to very low read depth in some genes/individuals, several genes showed highly variable PatRatios within the population. A small X-and Y-axis jitter was added to reduce overplotting effects. Genes shown in black were significant (FDR <0.1), while those in red did not pass this statistical threshold for significance. The figure is divided into three panels left, middle and right panel. Genes in the middle panel were found significant in LCL and/or WB and reported as putatively imprinted in GTEx (Baran et al. 2015); genes shown in the left panel were found significant in LCL and/or WB but not reported in GTEx; and genes shown in the right panel represent those reported as putatively imprinted in GTEx but were not identified as showing significant evidence of imprinting in either LCL or WB. Some genes in the right panel such as *DLK1*, *MEG9*, *THEGL*, *DIRAS3*, *PWRN1* and *NDN* show evidence of parental expression bias, but the limited number of informative samples meant we did not consider these in our formal analysis. For genes with multiple UGFs (Supplementary Table 2), we plot Paternal Ratios for the UGF with the most significant p-value.

**Table 1.**
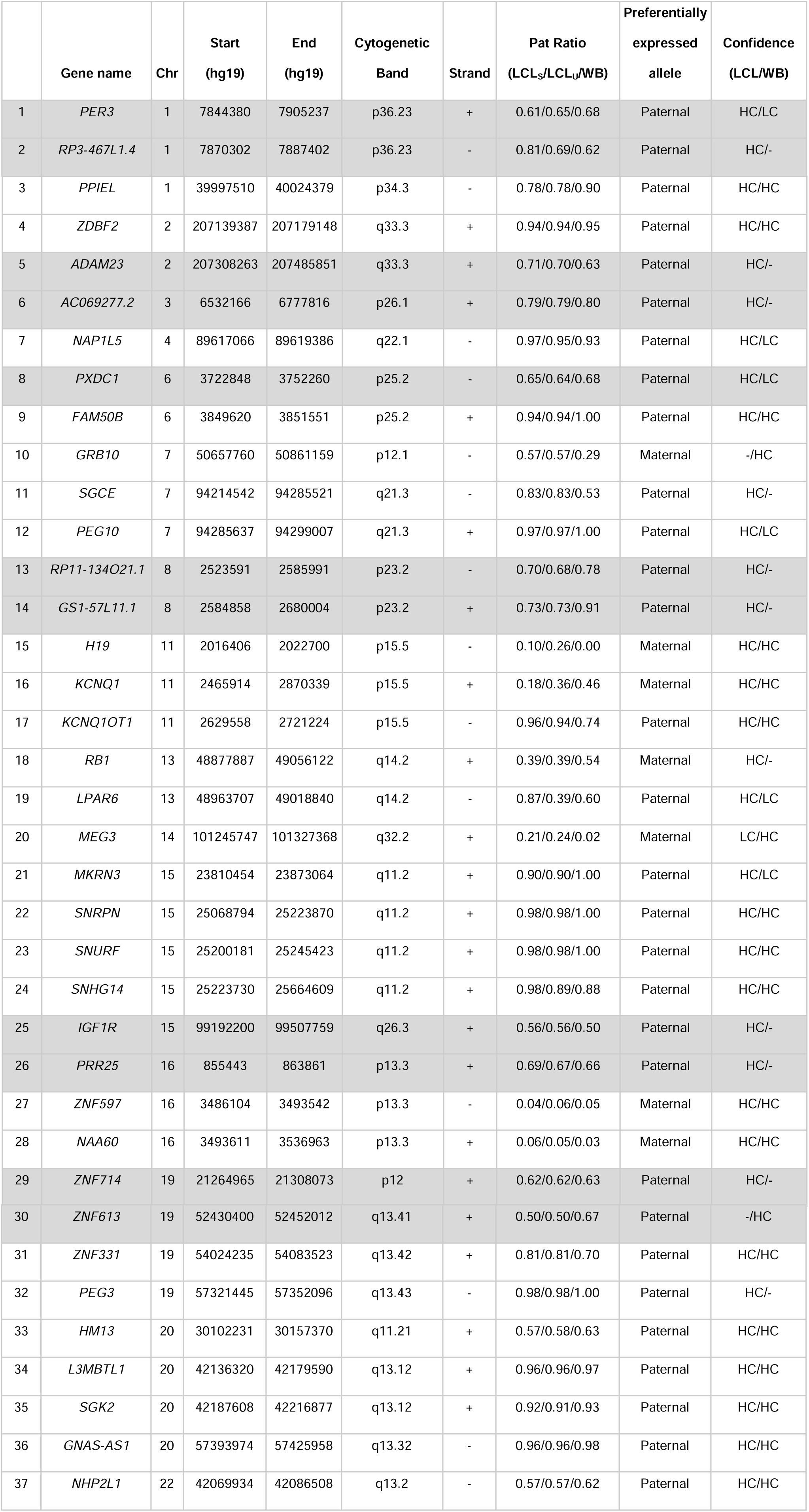
High confidence imprinted genes identified in LCLs and whole blood. High confidence imprinted genes were classified as those transcripts showing significant evidence of parental expression bias (at 10% FDR) by both statistical tests used in at least one of the two cohorts studied. Rows shaded grey indicate novel imprinted loci not reported in previous studies. For genes with multiple UGFs (Supplementary Table 2), we report Paternal Ratios for the UGF with the most significant p-value.

**Table 2.**
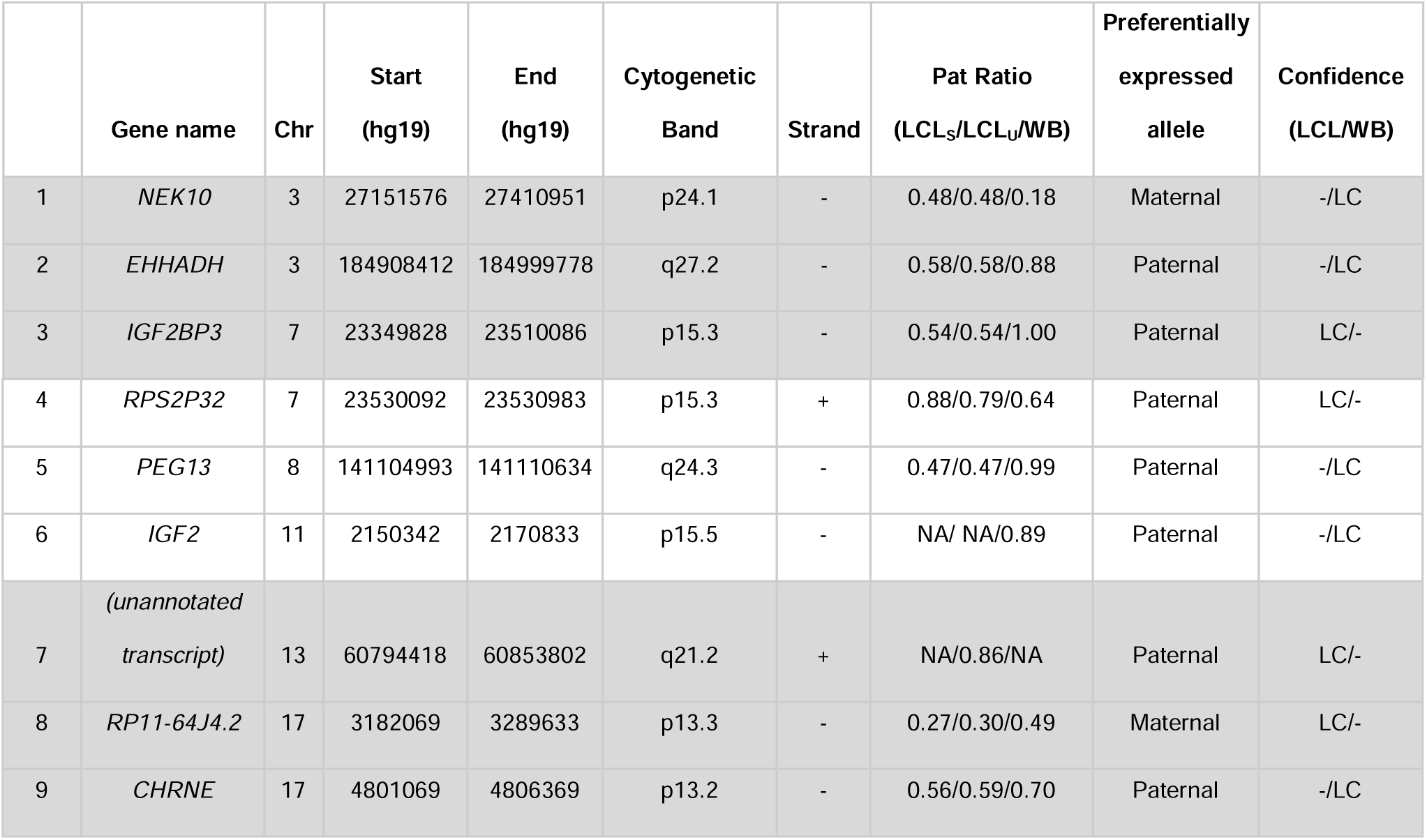
Low confidence imprinted genes identified in either LCLs or whole blood. Low confidence imprinted genes were classified as those transcripts showing significant evidence of parental expression bias (at 10% FDR) by just one statistical test in one of the two cohort studied. Rows shaded grey indicate novel imprinted loci not reported in previous studies. For genes with multiple UGFs (Supplementary Table 2), we report Paternal Ratios for the UGF with the most significant p-value.

For each dataset, we classified genes as high confidence if significant in both statistical tests (34 in LCLs and 20 in WB), and low confidence if a gene was identified as significant with only a single statistical test (4 in LCLs and 11 in WB). At 10% FDR using the Paired Sample Wilcoxon Signed Rank (WSR) test, we found 36 and 24 significant genes in LCLs and WB, respectively. With *ShrinkBayes* (SB), we found 37 and 27 significant genes in LCLs and WB, respectively at 10% FDR (Tables 1 and 2).

We compared the 45 imprinted genes in our dataset with those reported as imprinted by the Genotype-Tissue Expression (GTEx) project (Baran et al. 2015). Of the 29 genes previously reported as imprinted in either LCL or WB that were successfully assayed in our analysis, 19 showed significant parental expression bias in our study (Figure 2). In all cases we observed consistent directionality of parental bias between the two studies (Supplementary Table 3).

Using only female samples, we searched for signals of imprinting on the X chromosome. We first estimated X chromosome inactivation ratios (XCIR) in each female, removing those samples that showed highly biased XCIR (>80% silencing of one X chromosome), and then normalized allelic read counts for X-linked genes in each sample based on their XCIR. Analyses of these data resulted in one gene showing putative significant parental bias in LCLs (*RNA28S5*), and one gene in WB (*ARSD*). However, both were discounted as false positive signals due to clear reference bias in both cases (Supplementary Figure 1).

### Exclusion of potential confounders

It has been reported that LCLs can sometimes undergo clonal expansion, which in turn can lead to elevated rates of monoallelic expression (Proudhon and Bourc’his 2010). As this has the potential to create artifacts that might resemble imprinting, we utilized the XCIRs we defined in females to identify and exclude LCLs with possible clonality. Focusing only on those female LCLs without skewed XCIR (XCIRs between 0.2 and 0.8, n=45), we repeated the WSR test for imprinting on the 56 autosomal UGFs that had informative SNVs in at least 5 of these non-clonal LCLs. Even with this markedly reduced sample size, every gene tested showed very similar Paternal Ratios to those obtained in the full cohort of 165 LCLs, with 36 of the 38 (95%) genes that we report as being imprinted in LCLs achieving at least nominal significance for unequal expression of the two parental alleles (Supplementary Table 4). Thus we were able to exclude the possibility that artifacts due to clonality in the LCLs we studied were driving our results.

Other studies have indicated that DNA methylation can become altered during the transformation and extended culture of LCLs, raising the possibility that this might create artifacts in our LCL cohort. To assess the stability of DNA methylation at imprinted loci in LCLs, we compared published datasets of DNA methylation in LCLs and blood, and comparing these with methylation profiles in samples with genome-wide uniparental disomy that show loss of imprinting (Supplementary Figure 2). This analysis showed that there was no evidence for systematic loss of imprinting in LCLs, and that methylation at the differentially methylated regions of imprinted loci is broadly similar between blood and LCLs.

### Novel incompletely imprinted genes occur in clusters

Most previous studies have identified imprinted genes based on the complete silencing of one parental allele. However, our large population sample and the quantitative nature of our assay identified several genes with biallelic expression, but which showed a significant bias for increased expression of one of the two parental alleles (Figure 2). In many cases, these incompletely imprinted genes occurred in close proximity to previously known imprinted genes that show mono-allelic expression. For example, we identified *PXDC1*, which lies ~100kb distal to the known imprinted *FAM50B* at 6p25.2, as showing a 2:1 paternal expression bias (Figure 3). Similarly, *ADAM23*, which lies ~130kb distal to *ZDBF2* at 2q33.3, also exhibits ~2-fold over-expression from the paternal allele. Overall, we identified 11 clusters of imprinted genes (defined here as two or more imprinted genes separated by <500kb), with 25 of the 46 imprinted genes we report located in these clusters.

**Figure 3.**
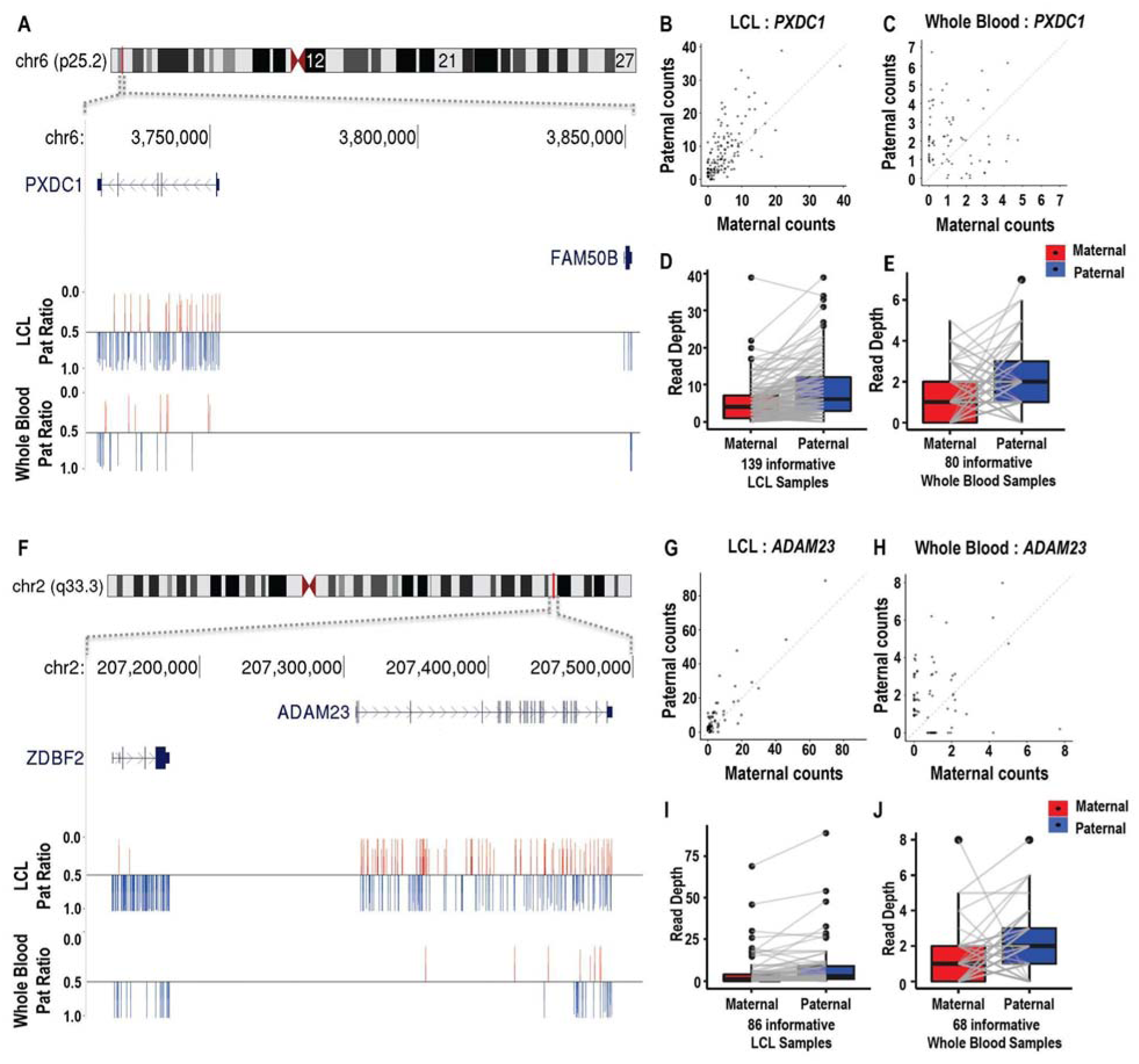
*PXDC1* and *ADAM23* are novel incompletely imprinted genes that lie adjacent to known imprinted genes. **(A-E)** *PXDC1* lies ~100kb distal to the known paternally expressed gene *FAM50B* at 6p25.2, and, although bi-allelically expressed, shows approximately 2-fold higher expression from the paternal allele in both LCLs (**B** and **D**) and WB (**C** and **E**). **(F-J)** *ADAM23* lies ~130kb distal to the known paternally expressed gene *ZDBF2* at 2q33.3, and also exhibits ~2-fold over-expression from the paternal allele in LCLs (**G** and **I**) and WB (**H** and **J**). In **(A)** and **(F)**, the mean fraction of reads transcribed from the paternal allele at every informative SNV position (the Pat ratio) is shown as bar, using a baseline of 0.5 (corresponding to equal expression of the two parental alleles). SNVs with preferential paternal expression (Pat ratio >0.5) are shown in blue, while SNVs with preferential maternal expression (Pat ratio <0.5) are shown in red. In **(D/E)** and **(I/J)**, vectors join the allelic expression values within each informative individual based on the sum of total RNA-Seq reads overlapping phased heterozygous SNVs within each gene.

To systematically investigate whether weaker imprinting localizes around strongly imprinted genes, we used data from a sliding window analysis across the genome in LCLs (detailed below) to test for an enrichment of parental expression bias around known imprinted genes. Here we divided the genome into 25kb bins, within each bin aggregated maternal and paternal read counts for all available heterozygous SNVs, and calculated the WSR p-value for parental expression bias for each 25kb window. We took the set of all 25kb non-overlapping windows that lie within ±250kb of strongly imprinted genes (those with Paternal Ratios ≤0.1 or ≥0.9), removing any windows that overlapped other strongly imprinted genes, and compared the p-values for parental expression bias in the resulting set of 175 25kb windows versus all 25kb windows in the rest of the genome (n=58,951). We observed that regions surrounding strongly imprinted genes are significantly enriched for signals of parental expression bias (permutation p=0.0005). Thus our observations extend the known clustering of imprinted genes in the mammalian genome, showing that effects of genomic imprinting can extend over broad regions, and cause genes to show differing extents of parentally biased expression.

In another example, we identified two anonymous transcripts *RP11-134O21.1* and *GS1-57L11.1* at 8p23.2 as novel imprinted genes showing a ~2:1 preferential expression of the paternal allele (Figure 4). Our previous studies of blood samples from patients with UPD (R. S.Joshi et al. 2016) identified a maternally methylated region located at the bidirectional promoter of these two transcripts, thus providing independent validation of our results.

**Figure 4.**
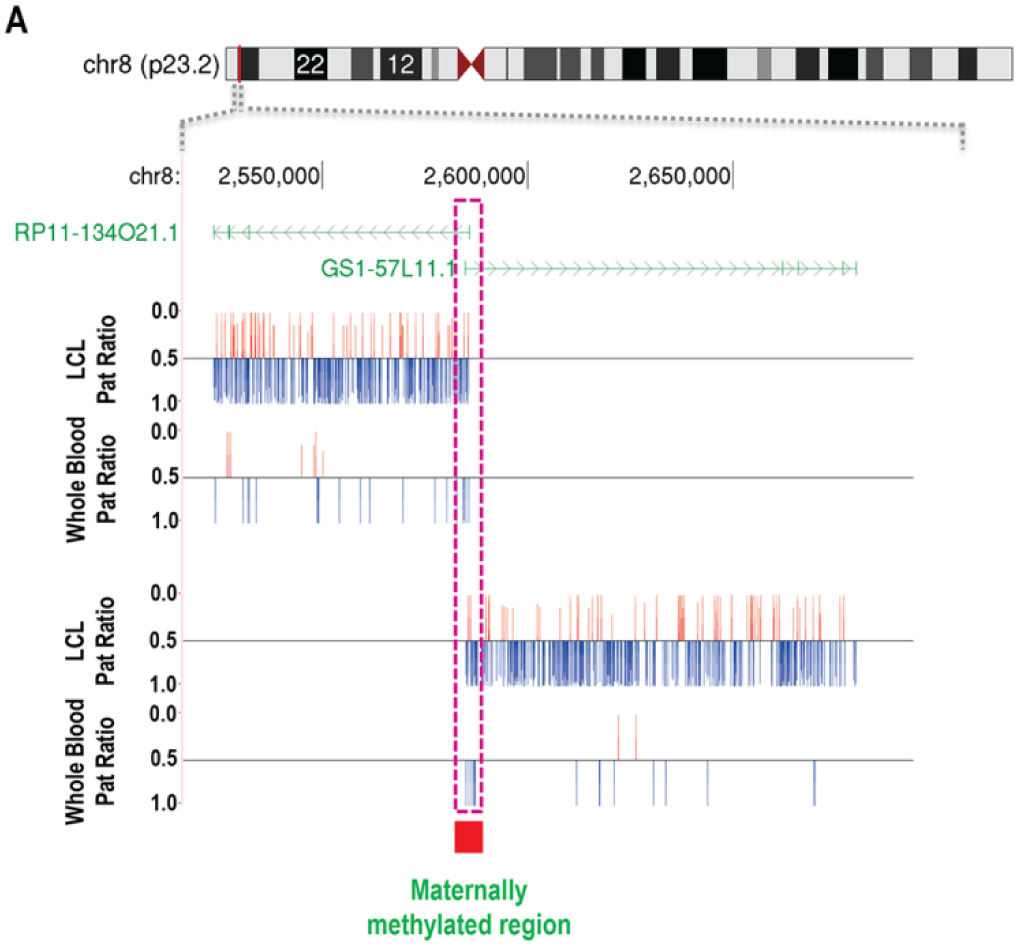
Two novel imprinted transcripts located at 8p23.2 share a bidirectional promoter that coincides with a maternally methylated locus. *RP11-134O21.1* and *GS1-57L11.1* lie in an antisense orientation, and both show ~2-fold expression from the paternal versus maternal allele in LCLs. Prior DNA methylation studies (R. S. Joshi et al. 2016) identified a region of increased maternal methylation located at the shared promoter of these two transcripts, confirming parent-of-origin effects at this locus, and indicating this as the likely regulatory element controlling imprinted expression at this locus.

### Strand specific RNA-Seq data reveals overlapping sense/anti-sense genes with opposite imprinting

In LCLs, the availability of strand-specific RNA-Seq data allowed the quantification of maternal and paternal counts from the forward and reverse strands separately. In the majority of cases, results obtained using stranded data were very similar to those obtained when aggregate data from both strands were considered. However, at loci where overlapping genes were transcribed from both forward and reverse strands, the use of unstranded data yielded misleading results. For *KCNQ1*/*KCNQ1OT1*, *RB1*/*LPAR6*, *NAA60*/*ZNF597,* and *PER3*/*RP3-467L1.4* only the use of strand-specific data was able to unambiguously determine the imprinting status of these genes (Figure 5). Notably, the strand-specific data demonstrated that several sense and antisense transcript pairs displayed opposite parental bias: *KCNQ1* is maternally expressed, whereas *KCNQ1OT1* is paternally expressed; *RB1* is maternally expressed, whereas *LPAR6* is paternally expressed (Figure 5 and Table 1).

**Figure 5.**
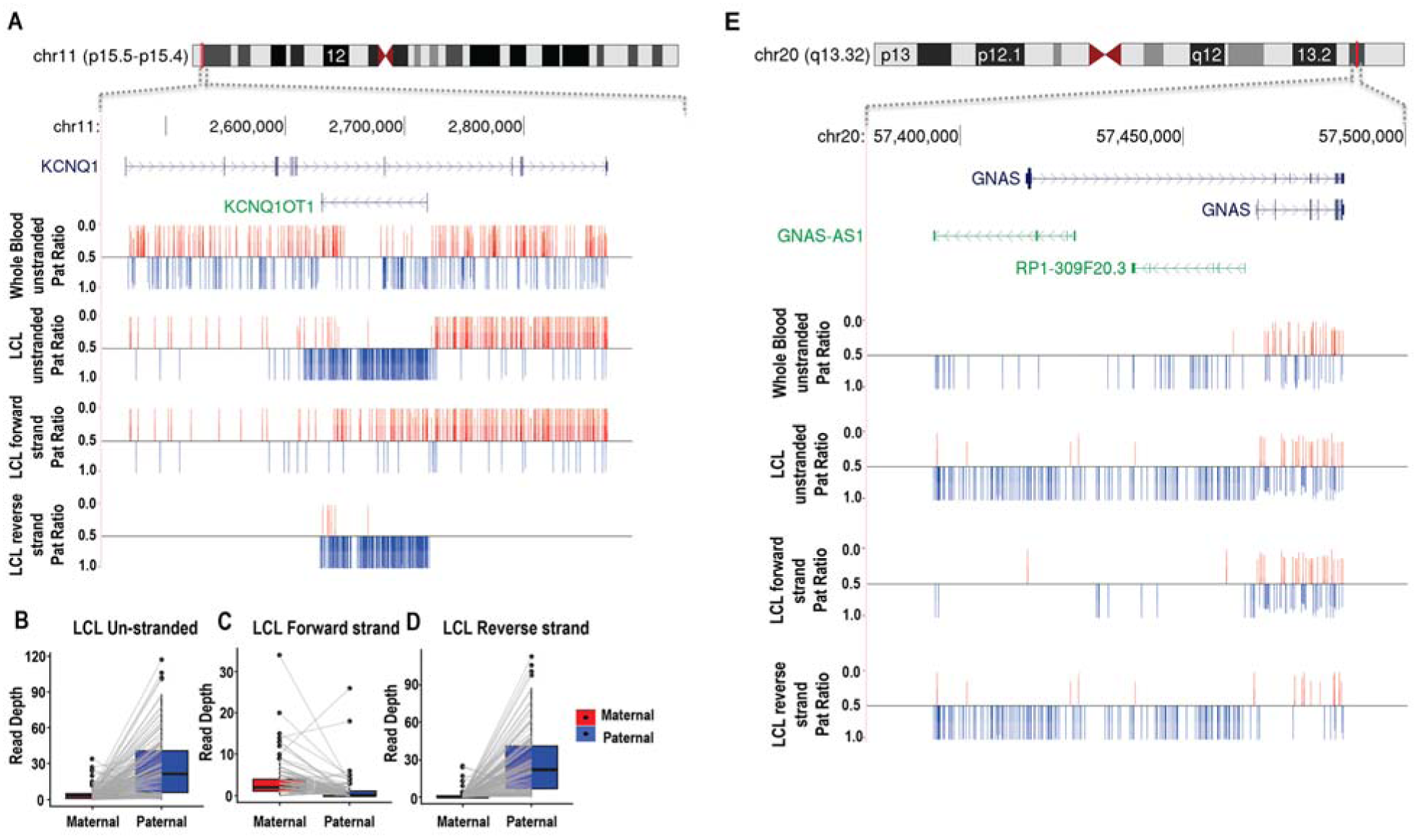
Stranded RNA-Seq data provides improved resolution of imprinting at overlapping antisense genes. Several loci in the genome contain multiple imprinted transcripts, including pairs of overlapping antisense genes with opposite imprinting patterns. Strand-specific RNA-Seq provided considerably improved ability to discern the correct imprinting patterns at these loci when compared to the use of unstranded libraries. **(A-D)** *KCNQ1* and *KCNQ1OT1* lie within the 11p15.5 imprinted region. *KCNQ1* on the plus strand is maternally expressed, while *KCNQ1OT1* on the negative strand is paternally expressed. In whole blood where only unstranded data was available, no significant parental bias was detected from either transcript, likely due to the combined signal from the two overlapping transcripts giving the appearance of biparental expression. However, the use of stranded RNA-Seq in LCLs clearly shows the that the two transcripts are antisense and have opposite imprinting patterns. **(E)** Similarly, *GNAS* and *GNAS-AS1* are antisense transcripts located in 20q13.32. In LCLs, the stranded RNA-Seq data shows that while *GNAS-AS1* is a paternally expressed imprinted gene, *GNAS* shows biparental expression.

### DIfferential imprinting at loci with multiple isoforms and overlapping transcripts

Previous studies have noted complex patterns of imprinting at certain genomic loci, such as isoform-specific imprinting, or imprinted genes that overlap with other non-imprinted genes (Court et al. 2014). Using data from the location of individual informative SNVs within the imprinted genes we report, we identified several loci that exhibited differential imprinting patterns among sub-regions of gene annotations.

One example of this phenomenon is *ZNF331*, which has multiple different isoforms with different transcription start sites. As shown in Figure 6, isoforms of *ZNF331* that start at the most proximal promoter show no evidence of imprinting, while other isoforms transcribed from more distal promoters show ~90% expression from the paternal allele. Previous reports (Court et al. 2014) have suggested that in blood leukocytes there is maternal-specific expression from the most proximal promoter of *ZNF331*, while our analysis indicates that in LCLs these isoforms show equal biparental expression. *HM13* shows a similar phenomenon of isoform-specific imprinting, with the longest isoforms showing a strong paternal expression bias, while shorter isoforms are biparentally expressed in blood (Supplementary Figure 3).

**Figure 6.**
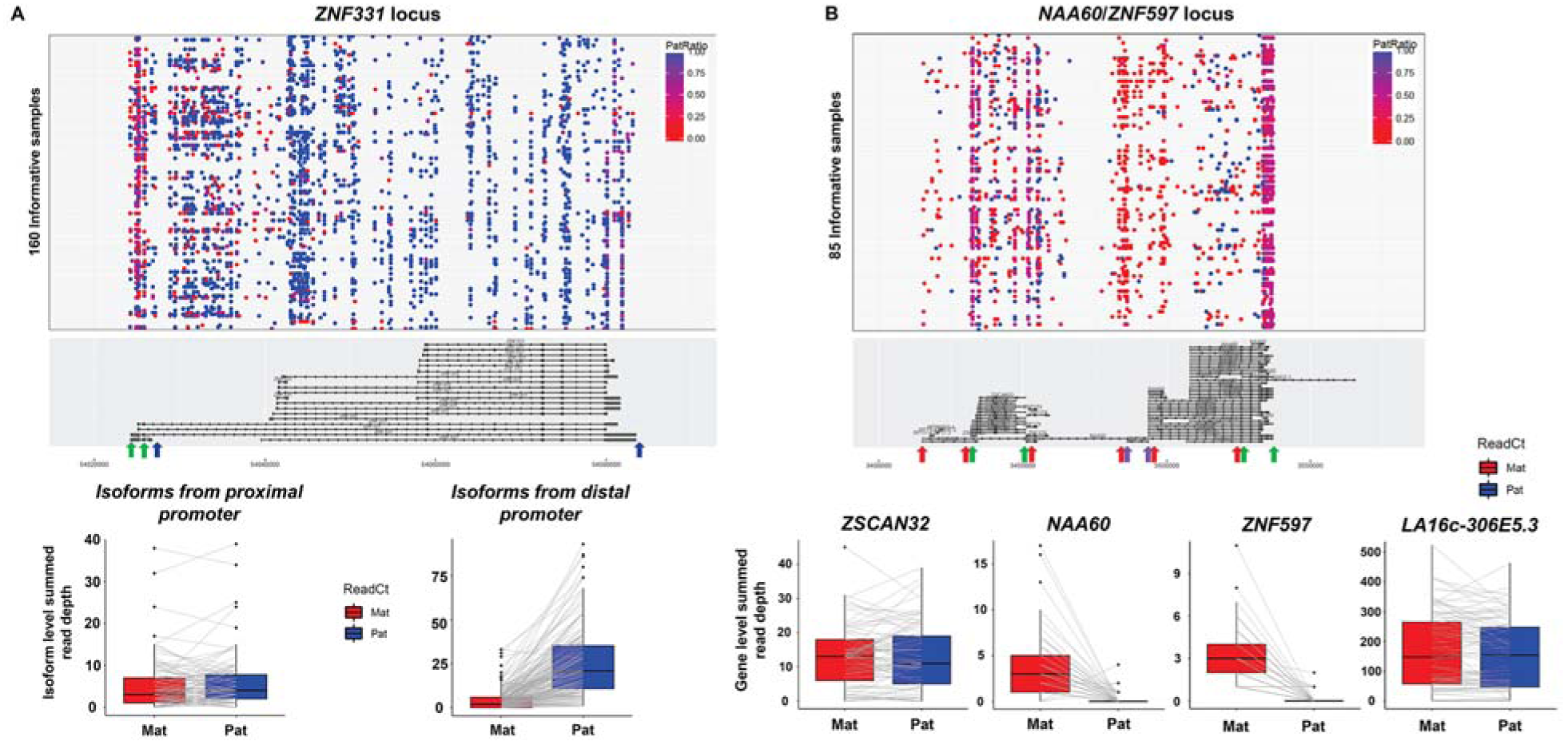
Complex patterns of imprinting at the *ZNF331* and *NAA60*/*ZNF597* loci revealed by phasing hundreds of transcribed SNVs. **(A)** Isoform-specific imprinting of *ZNF331* has been previously reported (Court et al, 2014, Stelzer et al, 2015) where the longer isoform has biallelic expression in human LCLs, while the shorter isoforms have paternal expression. Isoforms expressed from the proximal promoter (boundaries indicated by green arrows under gene plot) show biallelic expression (left boxplot), while longer isoforms of the gene (boundaries indicated by the blue arrows) show strong paternal expression bias (right boxplot). Thus, depending on the position of the observed heterozygous SNVs within the *ZNF331* gene, an individual may show different patterns of allelic bias. **(B)** Parental expression bias at *NAA60/ZNF597*, a complex locus that contains multiple overlapping imprinted and non-imprinted genes. The longest annotated isoform of *NAA60* overlaps the imprinted gene *ZNF597*, and also the bi-allelically expressed genes, *ZSCAN32*, *ZNF174* and *LA16c-306E5.3*. Considering SNVs within the boundaries of *ZSCAN32* and *LA16c-306E5.3* (regions defined by the green arrows) yields no evidence of imprinted expression, even though these are also contained within the longest annotated isoform of *NAA60*. However, considering SNVs located within the UGFs that uniquely describe *ZNF597* (defined by red arrows) or *NAA60* (defined by purple arrows) reveals almost exclusive expression from the maternal allele for these two genes. In the upper gene plots, each dot represents a single heterozygous SNV, which are colored to indicate the allelic ratio of the overlapping reads. Box plots show aggregate maternal and paternal read counts per individual.

The *NAA60/ZNF597* locus also shows similar complexity, containing multiple overlapping transcripts, only some of which are imprinted (Figure 6). *NAA60* has multiple isoforms, the longest of which overlaps several other genes, including *ZNF174*, *ZSCAN32* and *LA16c-306E5.3*. SNVs that overlap either *ZSCAN32*, *ZNF174* or *LA16c-306E5.3* show no evidence of parental bias, while SNVs that fall uniquely within *NAA60* or *ZNF597* show almost exclusive maternal expression.

Finally, careful inspection of the *TRAPPC9* locus enabled us to refine the signal of imprinting specifically to *PEG13*, which lies intronic within *TRAPPC9*. We observed a cluster of SNVs located in the center of the annotated *TRAPPC9* locus showing almost exclusive paternal expression, while SNVs located elsewhere in *TRAPPC9* showed equal expression from the maternal and paternal alleles (Supplementary Figure 3). Although the gene annotations we used (Gencode v16) includes multiple isoforms of *TRAPPC9*, none included exons that corresponded to the cluster of paternally expressed SNVs within *TRAPPC9*. Instead, the use of Refseq gene annotations included the 5.6kb transcript *PEG13* (*Paternally Expressed Gene 13*) that, like *TRAPPC9*, is expressed from the negative strand, and coincides perfectly with this cluster of paternally expressed SNVs that lie intronic within *TRAPPC9*. Thus, careful curation of this locus revealed that the imprinted signal we observed in blood comes solely from *PEG13*, and that the larger *TRAPPC9* gene is not imprinted in the cell types we studied.

### Genome-wide scan for imprinting outside of known gene annotations

In order to search for novel signatures of imprinting outside of current gene annotations, we utilized a sliding window approach to systematically analyze the entire genome in an unbiased fashion. We chose a window size of 25kb as this was close to the median transcript length, with a 5kb incremental slide. At each position, we aggregated maternal and paternal read counts for all available heterozygous SNVs within the 25kb window, and calculated the WSR test statistics (Supplementary Table 5). Using this approach, as expected, we identified significant associations at nearly all imprinted genes found using our gene-centric approach. In several cases (e.g. *ZNF331* and *ZDBF2*), significant signals of imprinted expression were observed downstream of annotated genes, which might represent transcriptional read-through beyond annotated 3’ boundaries (Supplementary Figure 4). However, we also identified a significant signal of expression outside of known gene annotations on 13q21.1 in the LCL population. Here, a cluster of 35 informative SNVs spread over ~8kb showed a strong paternal bias, with 87% of reads supporting transcription from the paternal allele in 73 informative samples. We propose that this represents a novel paternally imprinted transcript transcribed from the forward strand that apparently shares a bidirectional promoter with *LINC00434* (Figure 7). In support of this, data from the ENCODE Project in cell line GM12878 indicates the presence of an anonymous transcript at this position that is consistent in size and strand with our observations. There was no significant expression from this locus detected in whole blood.

**Figure 7.**
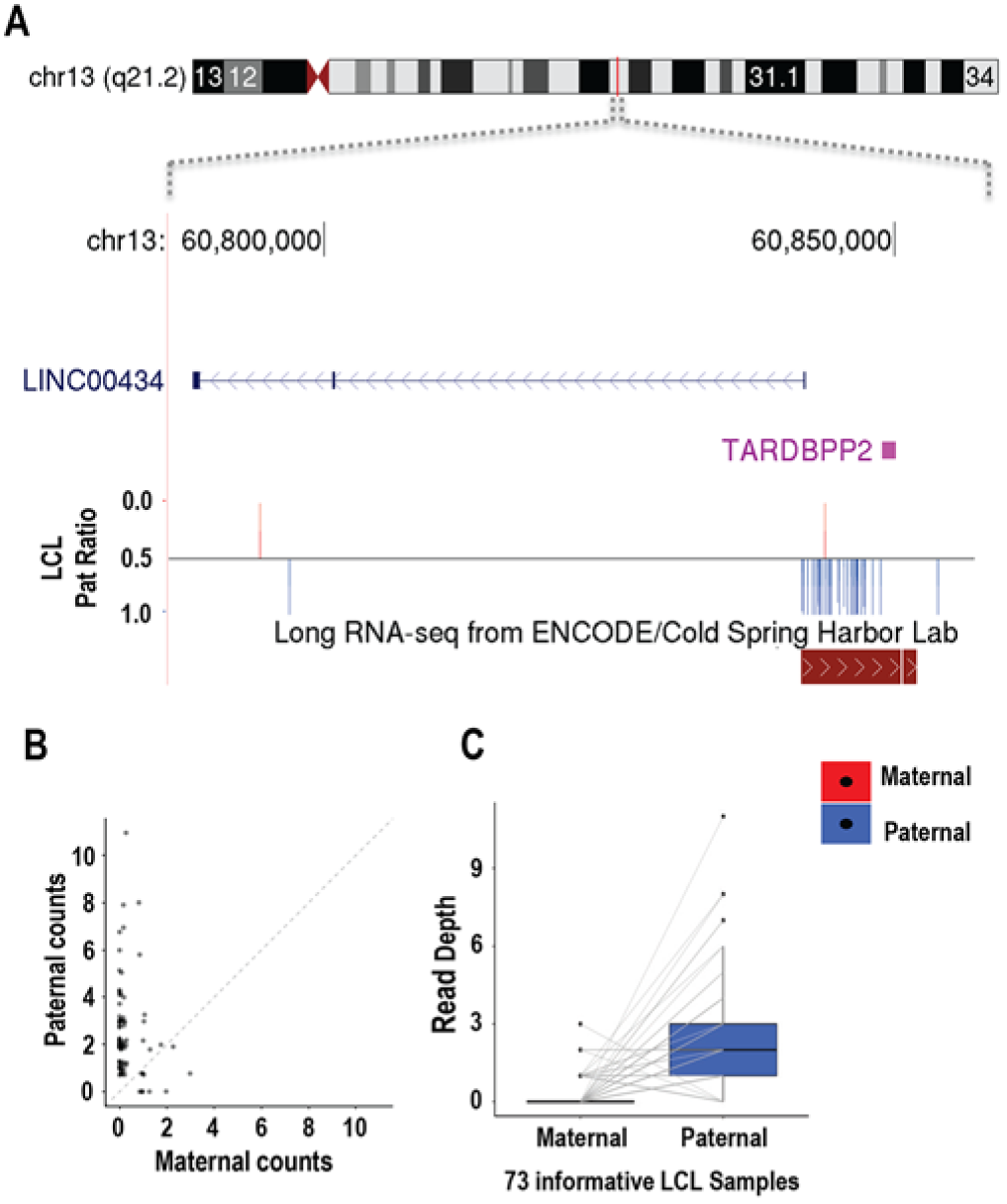
A novel putative imprinted lncRNA at 13q21.2. Using a sliding window analysis to interrogate the genome independent of gene annotations, we identified a cluster of 35 SNVs located in 13q21.2 (chr13:60,841,936-60,848,791, hg19) that showed a strong paternal expression bias. The putative transcript containing these SNVs is located on the forward strand, and apparently shares a bidirectional promoter with the non-coding RNA *LINC00434*. This SNV cluster overlaps a putative anonymous transcript identified in LCLs by the ENCODE project.

## Discussion

Here we report a detailed survey of imprinted gene expression in two human populations. We used a robust pipeline, incorporating the latest methods for allele-specific expression analysis, including rigorous removal of reads with potential mapping bias. Compared to previous methods, the availability of phased genotype information from whole genome sequencing of trios allowed direct assignment of expression levels from the two parental alleles at >2.8 million transcribed SNVs, providing a direct approach to assess imprinting genome-wide. This method provides a considerable increase in sensitivity compared to approaches where parental origin information is lacking, allowing us to detect much more subtle imprinting effects than have been observed previously.

Further, we developed a robust statistical framework to account for population heterogeneity of imprinting. While many previous studies have called events at the level of individual samples and variants, we studied nearly 300 independent trios, and employed two complementary statistical tests that considered aggregated read counts at the gene level. The paired WSR is a non-parametric test that has the advantage of a low false positive rate, but with reduced power at small sample size and low expression (Supplementary Figure 5). In contrast, SB uses the zero-inflated negative binomial distribution to fit the data, well-suited for zero-inflated count data such as RNA-Seq, providing increased power for genes with low expression. These approaches have the advantage of assessing differences between paternal and maternal RNA-Seq counts at multiple heterozygous loci across all individuals simultaneously, thus providing both increased robustness, and considerably greater power to resolve subtle biases in expression from the two parental alleles, when compared to the study of single data points. Consistent with prior studies, we found that utilizing aggregated read counts across all heterozygous sites per gene in each individual, including intronic reads and SNVs covered by only a single read, gave the most power in our analysis (Baran et al. 2015; DeVeale, van der Kooy, and Babak 2012). Finally, we filtered putative imprinted transcripts to remove false signals caused by reference bias, before manually curating each locus to resolve signals from overlapping and antisense transcripts. Importantly, curation to remove reference bias was an important step to avoid false positive imprinting signals: despite the fact that we masked non-unique genomic regions and applied stringent filtering to remove reads with ambiguous mapping, we still identified several genes with significant signals of parental expression bias that were attributable to reads mapping preferentially to the reference sequence (as assessed by statistical comparison of coverage of the reference and alternative alleles) (Supplementary Figure 1).

Overall, this pipeline led to the identification of 45 imprinted transcripts, 15 of which are novel, in addition to a novel unannotated imprinted locus on 13q21.2. Of the novel imprinted genes identified, two notable examples are *PER3* and *IGF2BP3*. *PER3* [Period, Drosophila, Homolog of, 3; OMIM# 603427] is a member of the Period family of genes and is expressed in a circadian pattern in multiple tissues (Zylka et al. 1998). *PER3* is one of several genes that regulate circadian rhythms, and has been linked to Seasonal Affective Disorder by both human and mouse studies (Delaunay et al. 2000; Zhang et al. 2016). *IGF2BP3* [Insulin-like Growth Factor 2 mRNA-Binding Protein 3; OMIM# 608259] binds to the 5’ UTR of the imprinted gene *IGF2*, suggesting it has a role in the regulation of *IGF2* production and is expressed ubiquitously across fetal and adult tissues (Monk et al. 2002; Nielsen et al. 1999). While previous reports have shown that *IGF2BP3* is bi-allelically expressed, we identify a slight bias for increased expression from the paternal allele. This may point at a coordinated PofO-based regulation of *IGF2* signalling cascade. Notably a maternally methylated CpG island associated with *RPS2P32* gene lies ~22kb upstream of *IGF2BP3* (R. S.Joshi et al. 2016).

Classical studies of imprinting typically define imprinted genes as showing monoallelic expression from just one of the two parental alleles. However, recent studies in mouse have identified examples of incomplete, or non-canonical, imprinting (Bonthuis et al. 2015) – such genes are bi-allelically expressed, but show a significant allelic bias, such that the two parental alleles are expressed at different levels. Our study also finds multiple examples of incomplete imprinting in the human genome, and we report nine imprinted genes that each show consistent 2-to 3-fold higher expression from the paternal allele. In several cases, these incompletely imprinted genes occur in close proximity to known imprinted genes that show mono-allelic expression, consistent with the known clustering of imprinted genes (Edwards and Ferguson-Smith 2007). While it is possible that some of these genes with incomplete imprinting in blood and/or LCLs might be fully imprinted (*i.e.* monoallelically expressed) in other tissues, we note that none were found in a prior survey of imprinting that assayed 34 human tissues (Baran et al. 2015), making this unlikely.

Of note, we observed that some genes showed large apparent variations in Paternal Ratios (Figure 2), and we found several different factors contributing to this phenomenon. In some cases, such as *PXDC1* or *PER3*, this was apparently due to stochastic variation as a result of low read depth. For example, where an individual has a single heterozygous SNV in a gene that is covered by only two RNAseq reads, the possible paternal expression ratios are 0, 0.5 or 1. Thus, in the case of a gene with low expression and incomplete imprinting, wide variations in the allelic ratios among different individuals will be observed as a result. In other cases, apparent variability of allelic ratios could be attributed to the fact that some genes showed isoform-specific imprinting patterns. For example, *ZNF331* has multiple different isoforms with different transcription start sites: in LCLs, those transcribed from the distal promoters show ~90% expression from the paternal allele, while isoforms transcribed from the most proximal promoter showed no evidence of imprinting. Thus, depending on the position of heterozygous SNVs within *ZNF331* carried by any one individual, the allelic ratio varied accordingly. Similar variability was also observed for *NAA60*, stemming from the fact that there are several overlapping annotated genes at this locus, all of which have much higher expression levels in LCLs than *NAA60*. As a result, the paternal ratio of any one SNV within *NAA60* is highly dependent upon its position within the locus. SNVs that overlap either *ZSCAN32, ZNF174* or *LA16c-306E5.3* showed no evidence of parental bias, while SNVs in regions that overlap only *NAA60* or *ZNF597* showed almost exclusive maternal expression (Figure 6).

In addition to a gene-centric approach, we also utilized a sliding window analysis to screen for imprinted transcription across the genome, independent of known transcript annotations. This identified a novel imprinted locus at 13q21.2, apparently corresponding to an anonymous lncRNA approximately 8kb in length. This imprinted transcript is antisense to *LINC00434*, with the two genes apparently sharing a bidirectional promoter. Although we did not detect any expression from *LINC00434* in LCLs, given that these two genes are likely transcribed from the same promoter, we hypothesize that *LINC00434* may also be imprinted.

Given a previous report of sex-specific variations in imprinting (Baran et al. 2015), we tested whether age or gender influenced the imprinting status for any of the 46 imprinted transcripts we identified. However, we did not detect any significant effects of these two variables on parental expression bias (Supplementary Note). Furthermore, as studies in mouse (Davies et al. 2005; Raefski and O’Neill 2005) have previously identified a cluster of imprinted genes on the X chromosome, and phenotypic studies in human have led to the suggestion that genes on the human X chromosome may also be subject to imprinting (Skuse et al. 1997), we specifically searched for imprinting on the X chromosome. Although this analysis utilized only female samples, and thus suffered a reduction in power compared to our analysis of the autosomes, we were unable to detect any evidence to support the presence of imprinted genes on the human X chromosome.

We compared the list of genes we detected as imprinted with those found in previous studies of imprinting (Baran et al. 2015), and overall found good concordance. However, for ten genes that were reported as imprinted in the GTEx cohort we did not observe evidence of imprinting, despite these genes having sufficient informative SNVs to be adequately assessed in our samples (*UTS2*, *MEST*, *UBE3A*, *PLAGL1*, *CPA4*, *MAGI2*, *INPP5F_V2*, *PRSS50*, *THEGL*, *RP11-7F17.7*). We note that of these ten genes, *MEST*, *UBE3A*, *PLAGL1*, *CPA4*, *MAGI2* and *INPP5F_V2* have all been reported as imprinted in other prior studies. While it is possible these may represent false-negatives in our analysis, many apparently show tissue-specific imprinting, with normal biparental expression in blood and LCLs, thus explaining our results (Kosaki et al. 2000; Vu and Hoffman 1997; Valleley, Cordery, and Bonthron 2007; Kayashima et al. 2003). In addition, we note that *UTS2* overlaps and is antisense to *PER3*, a gene which we identify as showing a weak paternal bias in LCLs. Given our improved methodology that utilized strand-specific RNA-seq, we suggest that the previously reported imprinting of *UTS2* instead likely reflects paternally-biased expression of *PER3*. Given the improved resolution of strand-specific over unstranded RNAseq data, we suggest that future expression-based studies of imprinting should utilize this approach where possible.

Our study has some limitations. Primarily, as our approach relies on measuring read depth over transcribed SNVs, we were limited to the study of genes that both contained heterozygous variants, and were expressed at sufficient levels to be analyzed. Thus, genes that were not expressed at detectable levels in a sufficient number of individuals, or which lacked heterozygous variants in our samples, were not assayed. Similarly, we had little discriminatory power to detect imprinting for genes that contained very few SNVs in our cohort, or for those that were expressed at very low levels. Further, as we studied samples of peripheral blood and LCLs, we were unable to detect genes that show imprinting confined to other tissues (Baran et al. 2015). Finally, as the LCLs we studied are immortalized cell lines, it is possible this process may have disrupted epigenetic processes such as imprinting. However, arguing against this possibility, there was both strong concordance of our results obtained in LCLs with previous studies of imprinting, and several of the novel imprinted genes detected in LCLs were also supported by methylation and/or RNA-Seq data from whole blood (Baran et al. 2015).

Given that our study assessed the imprinting status of ~41% of human transcripts, and identified 45 that are imprinted, our findings are broadly consistent with previous projections that have suggested that the human genome likely contains approximately 100 genes that are imprinted in somatic tissues (Barlow 1995).

## Methods

### Strand-specific RNA-Seq in 165 Lymphoblastoid Cell Lines

We generated RNA-Seq data from lymphoblastoid cell lines (LCL) for 57 CEPH (CEU), 58 Yoruba (YRI) and 50 Han Chinese (CHS) samples, all of whom were offspring of multi-generation pedigrees studied as part of The HapMap (http://hapmap.ncbi.nlm.nih.gov/) and/or 1000 Genomes (http://www.1000genomes.org/) Projects. Samples are listed in Supplementary Table 6.

#### Genotype data processing

For 163 samples, genotype data from the complete mother/father/child trio were available, while for the two samples, genotype data for only one parent was available. We obtained 1000 Genomes and HapMap project data from multiple releases: this included data from The 1000 Genomes Project Phase 1 and Phase 3 generated from low-coverage Illumina whole genome sequencing, high coverage Complete Genomics whole genome sequencing data, exome sequencing, Illumina Omni 2.5M SNV array data, and HapMap3 project data genotyped on Illumina 1.6M and Affymetrix 6.0 SNV arrays. We included high quality filtered and curated DNA genotype data from the final releases of all these resources and combined into population-specific datasets. We performed quality control on the merged data such as resolving strand inconsistencies, removing multi allelic SNVs, indels, removing SNVs not present in the 1000 genomes data and converting coordinates from hg18 to hg19 where required using PLINK (versions 1.07 and 1.9) (Purcell et al. 2007; Chang et al. 2015), vcftools (version 0.1.15) (Danecek et al. 2011) and Beagle utilities.

Due to the differing genotyping approaches and resulting SNV densities available across different individuals, we performed combined imputation and phasing to increase SNV density and infer the two parental haplotypes in each offspring with Beagle 4.0 (S. R. Browning and Browning 2007). This used family pedigree information with the 1000 Genomes Phase3 reference panel downloaded from Beagle website (http://bochet.gcc.biostat.washington.edu/beagle/1000_Genomes_phase3_v5). Using 493 HapMap samples from the CEU, YRI and CHS populations, we created population specific-reference panels to improve imputation accuracy. Since many of the samples in our target panel are also part of 1000 Genomes Project reference panel, for each population group we created subsets of target and reference panel in such a way that there are no overlapping samples in two sets, and imputed and phased each of these subsets of target panel separately. Each chromosome was divided into segments to efficiently perform imputation and phasing, and these segments were subsequently merged together to yield chromosome-wide imputed and phased genotypes. Imputed genotypes were filtered to retain only high-quality genotypes (R^2^≥0.95). We also removed sites with Mendelian errors in each trio, Hardy-Weinberg Equilibrium p<10^-4^, and retained only biallelic SNVs with Minor Allele Frequency ≥5% in at least one of the three ethnicities in the cohort. This yielded ~3.9 million high-quality SNVs phased for parental origin.

To reduce phase-switch errors introduced during phasing that would result in incorrect parental origin assignment of SNVs, we used an R script developed in-house (https://github.com/SharpLabMSSM/PofOAssignment). This method utilizes the phased genotypes generated using BEAGLE, as follows: Each offspring’s haplotype is compared with the parental haplotypes using a sliding window of 100 SNVs with 50 SNV incremental slide. Within each window we check for perfect matches between each offspring haplotype, and the four possible haplotypes within the parents. Parental origin assignments for each haplotype in the offspring are based on an unambiguous match to a single parental haplotype. This approach allows assignment of parental origin at uninformative sites where all members of the trio are heterozygous, and also provides an error check for phase switching. In the case when offspring’s haplotypes do not perfectly match a parental haplotype, the genotypes in the window are set to missing. Subsequently, we then recover any such lost sites using simple rules of Mendelian Inheritance to each individual SNV genotype in the trio. Thus, by using a combined approach leveraging both statistical phasing with rules of Mendelian inheritance, we are able to generate maximally informative assignment for parental origin at heterozygous SNVs, with a minimal error rate.

#### Sample preparation

Lymphoblastoid cell lines were obtained from the Coriell Institute (Camden, NJ). Cells were grown in RPMI1640 media supplemented with 1mM L-glutamine, 10% FBS and 100u/L each of penicillin and streptomycin, according to recommended protocols. Total RNA was extracted from frozen cell pellets (5-10 million cells) using TRIZOL, according to manufacturer’s instructions (ThermoFisher Scientific). Strand-specific RNA-Seq libraries were prepared using NEBNext Ultra Directional RNA Library Prep Kit from Illumina. 1μg of total RNA was used as input, polyA+ selected, followed by strand synthesis was performed. Libraries were sequenced on an Illumina Hiseq 2500 instrument, with 10 samples pooled per lane, to generate 100bp single-end reads to a median depth of ~16 million reads per sample.

#### RNA seq data processing

Quality control analysis was performed on RNA-Seq reads using fastqc (version 0.11.2) (http://www.bioinformatics.babraham.ac.uk/projects/fastqc). Over-represented sequences were removed using trimmomatic (version 0.32) (Bolger, Lohse, and Usadel 2014), and trimmed reads ≥30bp in length were kept. Cleaned reads were mapped to the human reference genome (hg19) with Gencode v16 annotations using the STAR aligner (version 2.3.0) (Dobin et al. 2013), yielding a mean of 79% uniquely mapped reads respectively. Picard (version 1.112) (https://github.com/broadinstitute/picard) was used for intermediate bam file processing such as add read groups, sorting and merging bam files of the same samples. To correct for mapping errors and biases which can result in false-positive allele-specific read-assignments, we used a collection of utilities in the WASP software (version 0.1) (van de Geijn et al. 2015), resulting in the removal of a mean of 36% of reads that overlapped SNVs in each sample, for which unambiguous allelic assignment could not be made. After parental-origin assignment for SNVs in each offspring, heterozygous sites were used to determine allele-specific expression. We first quantified reference and alternate RNA-Seq reads mapped at heterozygous loci using AlleleCounter (v0.2, https://github.com/secastel/allelecounter) implemented in Python (Castel et al. 2015). Then, reference and alternate allele counts were used with PofO information to assign counts to the maternal and paternal alleles at each heterozygous site. Reads that did not uniquely map, or had base quality ≤10, were discarded. To further reduce mapping errors we applied additional filters, removing heterozygous SNVs that: (i) had a mappability score <1 (based on the “CRG GEM Alignability of 50mers with no more than 2 mismatches” track, downloaded from UCSC genome browser), (ii) overlapped CNVs with MAF ≥5% identified in samples from the 1000 Genomes and HapMap Projects (ftp://ftp.1000genomes.ebi.ac.uk/vol1/withdrawn/phase3/integrated_sv_map/ and common CNVs (Conrad et al. 2010), (iii) Segmental Duplications, and (iv) Simple Repeats (both downloaded from “Variation and Repeats” track group of the UCSC genome browser). These filters resulted in the removal of 21% of heterozygous sites, leaving ~3.1 million sites for downstream analysis.

### Unstranded RNA-Seq in 131 Whole Blood Samples

The Genome-of-the-Netherlands (GoNL) project (Genome of the Netherlands Consortium 2014) performed whole genome sequencing of 250 family trios, a subset of which also had whole blood transcriptomes sequenced as part of the BBMRI-NL Biobank-based Integrative Omics Study (BIOS) (Zhernakova et al. 2017; Bonder et al. 2017). From these, we utilized data from 131 children with whole blood RNA-Seq data that passed all quality criteria and had genotypes concordant with those obtained by whole genome sequencing (listed in Supplementary Table 7). The individuals were participants from one of four biobanks: LifeLines-DEEP, The Leiden Longevity Study, Netherlands Twin Registry, and the Rotterdam Study.

#### Genotype data processing

DNA genotypes of 250 Dutch families were phased and imputed using BEAGLE (B. L.Browning and Yu 2009) and IMPUTE2. An integrated phase panel was constructed using SNV genotype likelihoods from the GATK:UnifiedGenotyper as input for BEAGLE, treating all samples as unrelated. SHAPEIT2 and MVNcall19 were then used along with trio information to phase the complete set of SNVs. Each haplotype transmitted to the offspring, and therefore allelic parental origin, was then obtained from the phased haplotypes (Genome of the Netherlands Consortium 2014).

#### Sample preparation

Total RNA from whole blood was treated using Ambion’s GLOBIN clear kit, and subsequently processed for sequencing using the Illumina Truseq version 2 library preparation kit. Paired-end 50bp reads were generated using an Illumina HiSeq 2000 instrument, pooling 10 samples per lane. Read sets per sample were generated using CASAVA, retaining only reads passing Illumina’s Chastity Filter for further processing. Data was generated by the Human Genotyping facility (HugeF) of ErasmusMC (The Netherlands, see URLs). Full details are described in (Zhernakova et al. 2017).

#### RNA-Seq data processing

Initial quality control was performed using FastQC (v0.10.1). Removal of adaptors was performed using Cutadapt (v1.1) (Martin 2011). Sickle (v1.2) (N. A. Joshi, Fass, and Others 2011) was used to trim low quality ends of the reads (minimum length 25, minimum quality 20). The reads were mapped with the STAR aligner (v2.3.125) (Dobin et al. 2013) to human reference genome hg19 masked at all single nucleotide variants with MAF>0.01 in GoNL samples. Full details are described in (Zhernakova et al. 2017). To reduce the influence of reference bias, we utilized WASP (version 0.1) (van de Geijn et al. 2015) to remove reads that aligned to different genomic positions after substituting the variant site. A summary of the influence of masking SNV positions in the reference and utilizing WASP to remove reads that show ambiguous mapping positions is shown in Supplementary Figure 6.

To obtain the parent-of-origin allelic counts, we first computed RNA-Seq reference and alternative counts using the GATK (v3.6-0-g89b7209) ASEReadCounter tool (McKenna et al. 2010). A script was then used to re-label the reference and alternative counts with parental origin based on the transmitted allele, leaving ~0.9 million heterozygous sites with paternal and maternal read counts for downstream analysis. A summary of the complete analytical pipeline is shown in Supplementary Figure 7.

#### Statistical analysis to identify imprinted expression

Since overlapping genes are common in the eukaryotic genome (Sanna, Li, and Zhang 2008), care must be taken when assigning reads to specific transcripts. To avoid misassignment of reads at SNVs located within overlapping transcripts, we compiled all genes from Gencode annotations into a model where we consider overlapping regions of different genes as a separate unit, termed “unique gene fragments” (UGFs) (Supplementary Figure 8). The resulting gene models comprised 79,452 UGFs, and were used for assigning each heterozygous SNV to specific genes.

To maximize statistical power for detecting PofO biased expression, we summed read counts for all SNVs within each UGF. We calculated the paternal allelic ratio (defined as the fraction of reads derived from the paternally-inherited allele) for each individual using aggregated read counts across all informative SNVs within each UGF. We used the paternal allelic ratio of each informative individual to calculate the mean paternal ratio per UGF.

To formally test for parental bias in expression of UGFs, we utilized two complementary statistical approaches. We chose (i) a frequentist non-parametric approach, the Paired Wilcoxon Signed Rank (WSR) test, and (ii) an empirical Bayes approach *ShrinkBayes (van de Wiel et al. 2014)*. ShrinkBayes computes a Bayesian False Discovery Rate (BFDR), and we applied Benjamini–Hochberg False Discovery Rate (FDR) correction to the results of the WSR test, considering those UGFs with FDR q<0.1 (10% FDR) as showing significant evidence of imprinting. In each cohort, we only considered results for those genes in which at least 10% of individuals had ≥1 read informative for parental origin. Based on the results of these two tests, we classified predicted imprinted genes into those with high confidence (identified as significant by both tests) and low confidence (significant by one of the two tests). WSR test is a paired difference non-parametric test. It assigns ranks to the paternal/maternal differences with 0: mean difference in pairs is symmetric around 0. The test is robust against outliers and has no distributional assumption. ShrinkBayes is an advanced statistical method specifically designed to handle zero-inflated count data allowing multi-parameter inference and modeling of random effects in a Bayesian setting. It relies on INLA (Rue, Martino, and Chopin 2009) for the parameter estimation per gene, while borrowing information across genes by empirical Bayes type shrinkage of parameters. It allows a spike-and-slab prior for the parameter of interest (*patmat: mean difference in pairs*) to test *H_0_.* Per UGF, we use a simple model with a single predictor parameter for imprinting (patmat) and a random effect parameter (indiv) to account for within individual variability.

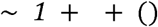

To assess the performance of the test procedures ShrinkBayes and WSR we developed a simulation scheme. ShrinkBayes is superior to WSR in terms of statistical power (Supplementary Figures 5 and 9) at a cost of increased computational resources. Using the two tests together reduces the false positive rate (Supplementary Figure 9), which motivates our definition of high-confidence genes.

Following statistical testing, we manually curated the UGF level results based on visual inspection of data plots, considering both gene annotations and strand-specific data in LCLs. Here we removed redundancies, and in the case of overlapping transcripts assigned imprinted expression to the correct gene. At several loci where we detected imprinted expression, gene annotations included transcripts with anonymous clone IDs. An example of this is the *L3MBTL1*/*SGK2* locus on chromosome 20. Here Gencode annotations include a transcript *RP1-138B7.5*, which is almost identical to an isoform of *SGK2*. In such cases, even though the transcript *RP1-138B7.5* was included in our initial list of significant imprinted genes, to avoid artificially inflating the number of imprinted transcripts we report, where these anonymous clone IDs likely corresponded to other annotated genes, we did not report them in our final curated list (Tables 1 and 2). Furthermore, although we filtered reads for potential mapping bias using WASP, we performed an additional check of UGF-level data for reference bias. We aggregated reference and alternate allele read counts at the UGF level, and applied a two-sided WSR test to check whether the distribution of reference and alternate read counts were significantly different after multiple testing corrections (5% FDR), removing genes that showed significant reference bias.

## Data Access

The raw and processed RNA seq data for 165 LCL samples have been deposited in the NCBI GEO database under the accession number GSE92521. The 131 WB STAR aligned BAM files (freeze 2) are submitted to European Genome-phenome Archive (EGA) under the study EGAS00001001077 and dataset accession number EGAD00001003937. The phased/imputed SNV data are part of the The Genome of the Netherlands (GoNL) Project with EGA accession number EGAS00001000644.

## Acknowledgements

We would like to thank Tuuli Lappalainen and Stephane Castel for facilitating the collaboration underlying this study. This work was supported by NIH grant HG006696 to AJS. Research reported in this paper was supported by the Office of Research Infrastructure of the National Institutes of Health under award number S10OD018522. The content is solely the responsibility of the authors and does not necessarily represent the official views of the National Institutes of Health. This work was supported in part through the computational resources and staff expertise provided by Scientific Computing at the Icahn School of Medicine at Mount Sinai. Work on the WB samples was performed within the framework of the Biobank-Based Integrative Omics Studies (BIOS) Consortium and the GoNL Project which are funded by BBMRI-NL, a research infrastructure financed by the Netherlands Organization for Scientific Research (NWO project 184.021.007).

## Author contributions

AJS and PACH conceived and planned the study. BJ, RM, KKG, SMK, HHMD, and MAW performed bioinformatic analyses. DH grew cell lines and prepared RNAseq libraries. BJ, RM, PACH, AJS and SMK prepared the manuscript.

## List of Supplementary Tables

**Supplementary Table 1.** All Unique Gene Fragments with FDR q<0.1, prior to manual curation.

**Supplementary Table 2.** Data for all Unique Gene Fragments in genome.

**Supplementary Table 3.** Comparison of imprinted genes detected in the current study with those reported by Baran et al. in GTEx samples.

**Supplementary Table 4.** Reanalysis of Unique Gene Fragments with FDR q<0.1 using only 45 non-clonal female LCLs without skewed X chromosome inactivation ratios.

**Supplementary Table 5.** Data for 25kb sliding windows with FDR q<0.1.

**Supplementary Table 6.** 165 LCLs used for RNAseq analysis, and their parents.

**Supplementary Table 7.** 131 whole blood samples used for RNAseq analysis.

## List of Supplementary Figures

**Supplementary Figure 1.** Reference bias can cause false positive signals of imprinting.

**Supplementary Figure 2.** DNA methylation at imprinted loci is conserved in both LCLs and blood, and does not show frequent loss of imprinting.

**Supplementary Figure 3.** Careful data curation resolves complex patterns of imprinting at the *HM13* and *TRAPPC9*/*PEG13* loci.

**Supplementary Figure 4.** Significant signals of imprinted expression extend downstream of known imprinted genes.

**Supplementary Figure 5.** Putative imprinted UGFs identified by ShrinkBayes and/or the paired Wilcoxon Signed Rank test as a function of underlying sample size (L) and mean expression (R).

**Supplementary Figure 6.** The effect of masking SNV positions and utilizing WASP on reference genome mapping bias.

**Supplementary Figure 7.** A summary of the analytical pipeline used for identifying parental bias in gene expression in whole blood samples.

**Supplementary Figure 8.** Definition of unique gene fragments (UGFs).

**Supplementary Figure 9.** Power estimates for ShrinkBayes and the paired Wilcoxon Signed Rank test on number of individuals and genes.

